# Chromatin tethering to the nuclear envelope by nuclear actin filaments: a novel role of the actin cytoskeleton in the *Xenopus* blastula

**DOI:** 10.1101/105593

**Authors:** Haruka Oda, Natsuki Shirai, Naoko Ura, Keita Ohsumi, Mari Iwabuchi

## Abstract

The *Xenopus* oocyte is known to accumulate filamentous or F-actin in the nucleus, but it is currently unknown whether F-actin also accumulates in embryo nuclei. Using fluorescence-labeled actin reporters, we examined the actin distribution in *Xenopus* embryonic cells and found that F-actin accumulates in nuclei during the blastula stage but not during the gastrula stage. To further investigate nuclear F-actin, we devised a *Xenopus* egg extract that reproduces the formation of nuclei in which F-actin accumulates. Using this extract, we found that F-actin accumulates primarily at the sub-nuclear membranous region and is essential to maintain chromatin binding to the nuclear envelope in well-developed nuclei. We also provide evidence that nuclear F-actin increases the structural stability of nuclei and contributes to chromosome alignment on the mitotic spindle at the following M phase. These results suggest the physiological importance of nuclear F-actin accumulation in rapidly dividing, large *Xenopus* blastula cells.

## Introduction

Filamentous or F-actin is one of the major cytoskeletal filaments in eukaryotes. It is the most flexible filament and can be rapidly depolymerized and repolymerized (Blanchoin *et al.* 2014). In collaboration with actin-binding proteins,F-actin is organized into various architectures, bundles, networks, and gels, the contraction of which can be induced. Because of the versatile nature of F-actin, the actin cytoskeleton is involved in a variety of cellular phenomena, particularly cellular morphogenesis and movement. The roles of F-actin in the cell, especially in the cortex, have been extensively investigated (Bezanilla *et al.* 2015). However, relatively few studies have addressed F-actin in the nucleus because nuclear actin is usually monomeric; accordingly, our understanding of the role of F-actin in the nucleus has been limited toa few cases (Huet *et al.* 2012; Grosse & Vartiainen, 2013). For instance, in serum-stimulated cells,nuclear F-actin transiently emerges and is involved in transcription activation via depletion of actin monomers, which suppress transcription factors through direct binding in the nucleus (Baarlink *et al.* 2013). It has been suggested that the transient assembly of nuclear F-actin is accelerated by the nuclear import of actin nucleators (Baarlink *et al.* 2013; Belin *et al.* 2015). A recent study demonstrated that DNA damage induces nuclear F-actin assembly through the actin nucleators Formin-2 and Spire-1/2 and that nuclear F-actin is involved in the repair of damaged DNA (Belin *et al.* 2015). Although these studies have shed light on the involvement of nuclear F-actin in chromatin regulation, the function of nuclear F-actin as a nucleoskeleton in these cases is not yet clear.

The nucleoskeletal function of F-actin has been demonstrated in the oocyte of the amphibian *Xenopus laevis* and in starfish. Early studies showed that the quiescent oocyte of *Xenopus* accumulates actin at high concentrations in its huge nucleus, the germinal vesicle (GV) (Clark & Merriam, 1977; Clark & Rosenbaum, 1979), in which the actin forms filaments (Kiseleva *et al.* 2004; Bohnsack *et al.* 2006). The accumulation of F-actin in the GV increases the structural stability of the large nucleus (Bohnsack *et al.* 2006) and, by forming an elastic scaffold, prevents nuclear particles such as nucleoli and histone locus bodies from gravitational sedimentation (Feric & Brangwynne, 2013). In starfish oocytes, actin mesh transiently forms in the nuclear region at the onset of the first meiosis, delivering chromosomes to the meiotic spindle through the contraction of the mesh (Lénárt *et al.* 2005). These studies reveal that F-actin functions as the nucleoskeleton in the oocyte.

Study of the *Xenopus* oocyte also revealed that there is a regulatory mechanism specific to the oocyte for the nuclear accumulation of F-actin. Nuclear transport of actin monomers is regulated by importin 9 and exportin 6 (Exp6), the nuclear import and export receptors, respectively (Stüven *et al.* 2003; Dopie *et al.* 2012). It has been demonstrated that the Exp6 protein is not expressed in the oocyte, and, due to its absence, actin monomers accumulate in the GV, accelerating actin polymerization. Exp6 begins to accumulate during oocyte maturation and further increases after fertilization (Bohnsack *et al.* 2006), suggesting that F-actin is unlikely to accumulate in the nucleus in *Xenopus* embryonic cells. However, another study showed that fluorochrome-labeled actin monomers accumulate in the nucleus and exhibit A network-like pattern in *Xenopus* egg extracts (Krauss *et al.* 2003), suggesting nuclear F-actin assembly in eggs. Thus, it remains to be determined whether F-actin accumulates in the nucleus in *Xenopus* embryos. Following fertilization, *Xenopus* embryos undergo cleavage, progressing through rapid cell cycles with no growth in which nuclear assembly and disassembly alternate in ~15-min intervals. Thus, it would be interesting to know whether F-actin is involved in the dynamic changes of the nuclei in early *Xenopus* embryos.

In the present study, we examined the distribution of actin in early *Xenopus* embryonic cells and found that F-actin accumulates in blastula nuclei but not in gastrula nuclei. To investigate the role of blastula-specific nuclear F-actin, we devised a cell-free extract from *Xenopus* eggs that reproduces nuclear F-actin accumulation. Using this extract, we found that nuclear F-actin is involved in the regulation of chromatin binding to the nuclear envelope. Our results also indicate that nuclear F-actin contributes to stiffening of the nuclear lamina and facilitates chromosome alignment on the spindle. Finally, we discuss the physiological relevance of the actin nucleoskeleton in the *Xenopus* blastula.

## Results

### F-actin accumulates in the nucleus during the blastula stage

To examine whether F-actin accumulates in the nuclei of early *Xenopus laevis* embryos, particularly blastulae, we injected the mRNA of EGFP-tagged F-actin binding protein (EGFP-UtrCH or Lifeact-2EGFP) into fertilized eggs. When mRNA-injected embryos reached the late blastula stage (stage 9), they were mounted on a glass slide with buffer containinga DNA-staining fluorescent dye and gently squashed by adding a coverslip, followed immediately by observation through a fluorescence microscope. We found EGFP signal not only in the periphery, corresponding to the cortex, of mitotic blastomeres but also in the nuclei of interphase blastomeres (Fig. 1A, B, blastula). We also noted that masses of the EGFP-UtrCH signal were often associated with the mitotic spindle (Fig. 1A). When embryos at the gastrula stage (stage 10 or 11) were similarly observed, EGFP signal was present in the cell cortex but not in the nucleus (Fig. 1A, B, gastrula). These observations suggest that F-actin accumulates in the nucleus during the blastula stage but not after the gastrula stage. To confirm the nuclear accumulation of F-actin in blastulae, nuclei were isolated from embryos at various developmental stages, fixed immediately, and stained with fluorochrome-labeled phalloidin. Observations demonstrated that the phalloidin signal, an indicator of F-actin, was consistently detected in nuclei until the late blastula stage (stage 9), and the nuclear signal was markedly diminished at the early gastrula stage (stage 10) (Fig. 1C). Measurements of the nuclear size and the density of the nuclear phalloidin signal revealed that a high density of F-actin was present in the nucleus during the blastula stage (up to stage 9) while the nuclear size was progressively diminishing (Fig. 1D), as reported previously (Jevtić & Levy, 2015). Interestingly, when embryos reached the gastrula stage, the nuclear F-actin density abruptly decreased to a low level, while nuclear size remained unchanged (Fig. 1D).These results indicate that the nuclear accumulation of F-actin is specific to blastulae in *Xenopus* embryonic development.

**Figure 1.**
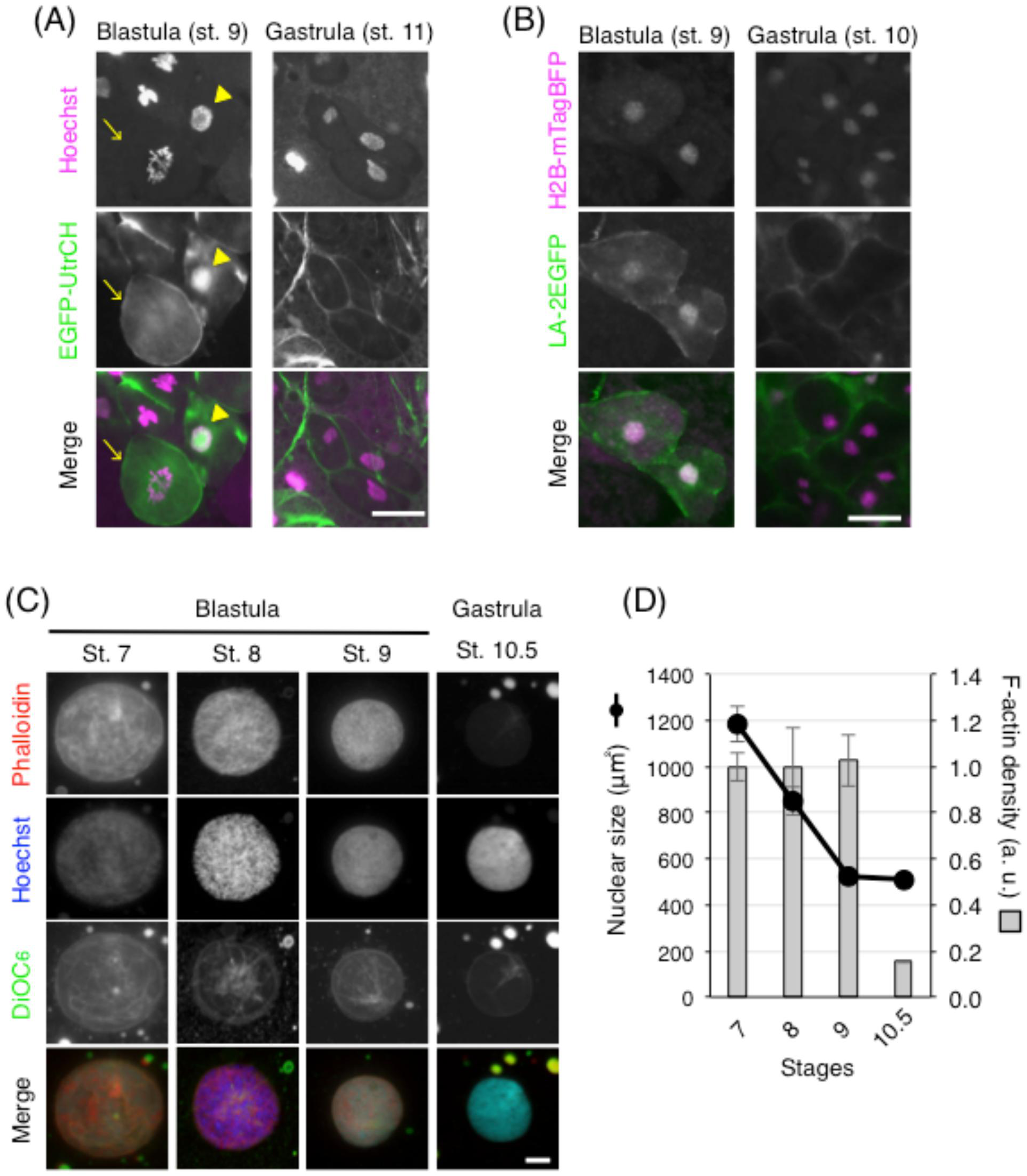
F-actin accumulates in the nucleus during the blastula stage. (A and B) *Xenopus* embryos were injected with EGFP-UtrCH (A) or LA-2EGFP and H2B-mTagBFP (B) mRNA and mounted on glass slides with Hoechst 33342 (A) or buffer (B) at the indicated stages before being gently squashed by a coverslip, followed immediately by observation by wide-field fluorescence microscopy. A nucleus accumulating EGFP-UtrCH and a mitotic blastomere are marked with an arrowhead and arrow, respectively (A). Scale bar, 50 m. (C) Nuclei were isolated from embryos at the indicated stages as described in Experimental Procedures. After fixation and staining of DNA, the nuclear envelope, and F-actin with DY-590-phalloidin, Hoechst 33342, and DiOC_6_, respectively, nuclei were observed by wide-field fluorescent microscopy. Representative images of each stage are shown. Scale bar, 10 m. (D) Changes in nuclear size and F-actin density of embryo nuclei. Means for nuclear size and F-actin density were obtained by measuring the nuclear area stained with DiOC_6_ and the intensity of nuclear DY-590-phalloidin signal, respectively, on the nuclear images taken as described in (C) with original threshold using NIS-element BR software (Nikon). Error bars represent SE (n > 20).

### F-actin accumulates in nuclei formed in egg extracts

To investigate blastula-specific nuclear F-actin in detail, we used cell-free extracts from *Xenopus* eggs, in which the nuclear formation of sperm chromatin is reproduced (Murray, 1991). Although *Xenopus* egg extracts have been conventionally prepared using an extraction buffer containing actin inhibitors (AI) to facilitate centrifugal separation of the cytoplasm, they have also been prepared without AI to study actin dynamics (Theriot *et al.* 1994; Ma *et al.* 1998; Sider *et al.* 1999; Valentine *et al.* 2005; Field *et al.* 2014; Abu Shah *et al.* 2015). Nuclear formation in such AI-free extracts, however, has not yet been reported. When the egg cytoplasm is separated by centrifugation from other cellular components in the absence of AI, a membranous layer containing pigment granules is formed above a semitransparent cytoplasmic layer, with a fuzzy border between the two (Fig. 2A, left; Field *et al.* 2014). In the present study, the pigmented and semitransparent layers were removed together and homogenized by pipetting before use to induce nuclear formation. When permeabilized sperm were incubated in the egg extract that had been prepared without the use of any inhibitors, they formed well-grown nuclei that resembled those formed in egg extracts containing AIs (cytochalasin B, CB or latrunculin A, LatA; Figs 2B, C, 3A). To examine whether F-actin accumulates in the nuclei, we induced nuclear formation in an AI-free extract that had been supplemented with EGFP-UtrCH protein. The result showed that the actin reporter accumulated in nuclei formed in the AI-free extract but not in those formed in AI-containing extracts (Fig. 2C, D). To confirm the nuclear accumulation of F-actin, nuclei formed in egg extracts were fixed on glass slides for staining with fluorochrome-labeled phalloidin. Observations of the nuclei demonstrated that the phalloidin signal was evident in the nuclei formed in AI-free extract, whereas no signal was detected in those formed in AI-containing extract (Fig. 3A). The egg extract that allows for actin polymerization in the nucleus is hereafter referred to as inhibitor-free extract (IFE) to distinguish it from the egg extract that contains AIs (inhibitor-containing extract, ICE).

**Figure 2.**
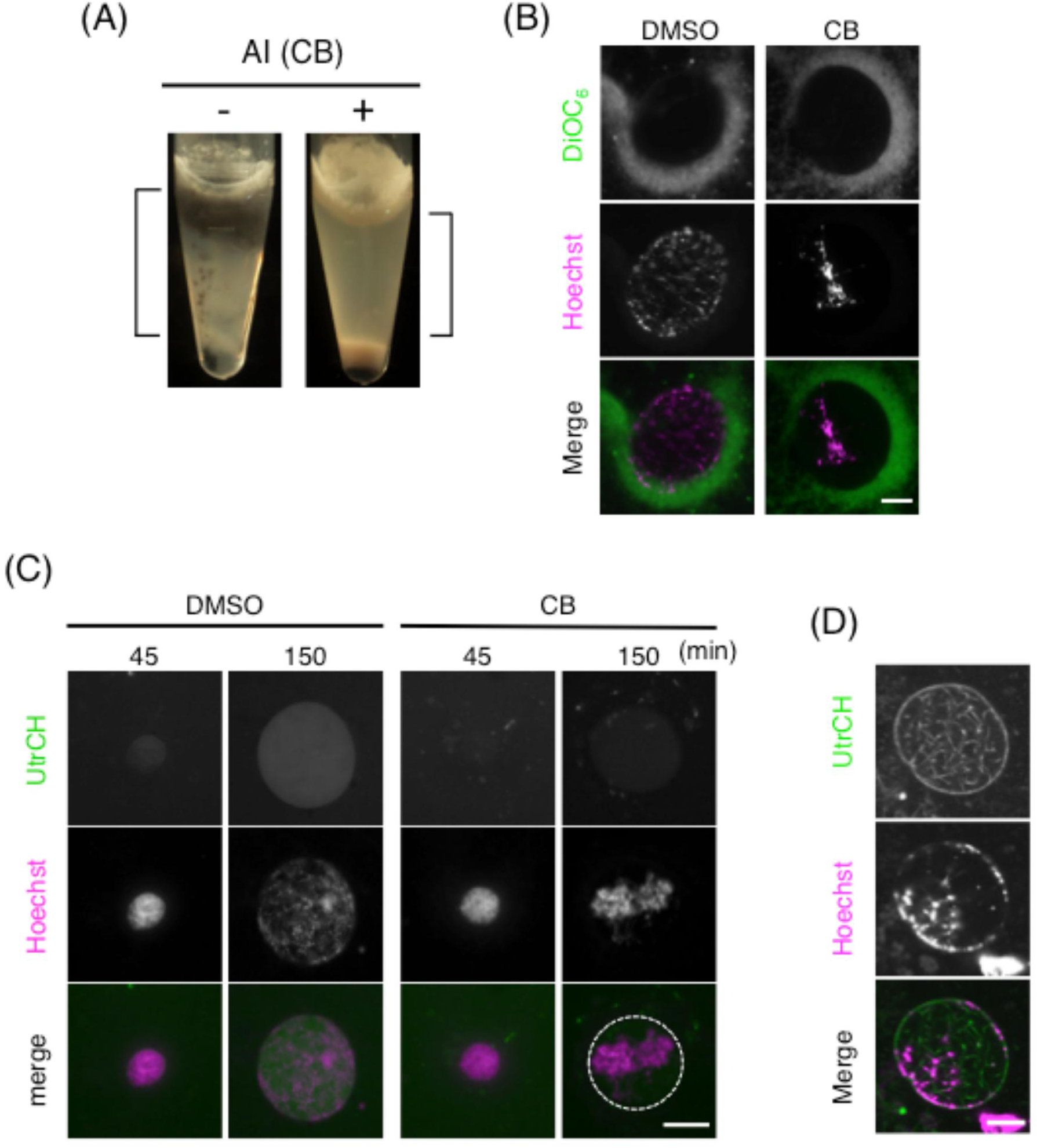
EGFP-UtrCH accumulates in nuclei formed in inhibitor-free egg extracts. (A)Centrifugal separation of the cytoplasm of *Xenopus laevis* eggs using cytochalasin B (CB)-containing (+) or CB-free (-) extraction buffer. Unfertilized *Xenopus* eggs were packed into a tube with buffer, and centrifugal separation was repeated twice as described in Experimental Procedures. In the absence of CB (-), a membranous layer containing pigment granules was formed beneath the lipid cap without clearly separating from the semitransparent cytoplasmic layer below. In the presence of CB (+), the semitransparent cytoplasmic layer was clearly separated from the lipid and membranous precipitates containing pigment gres. The cytoplasmic fractions between the lipid and precipitated membranes, which are indicated by square brackets, were removed and homogenized by pipetting for use as inhibitor-free (IFE) (-) and inhibitor-containing (ICE) (+) extracts. (B and C) AI-free (DMSO) and AI-containing (CB: cytochalasin B, 10 g/ml) extracts were supplemented with permeabilized sperm (B) or those along with EGFP-UtrCH protein (C) and were activated by the addition of CaCl_2_. Nuclei at 3 hours (B) or indicated time points (C) after incubation were observed by wide-field fluorescence microscopy with DNA (B, C) and the nuclear envelope (B) stained with Hoechst 33342 and DiOC_6_, respectively. The broken line in (C) indicates the nuclear contour. Scale bars, 20 m. Permeabilized sperm were added to IFE supplemented with EGFP-UtrCH protein and incubated for 120 min to induce nuclear formation. After Hoechst 33342 was added to the IFE, nuclei were observed without fixation by confocal microscopy as described in Experimental Procedures. Scale bar, 10 m.

**Figure 3.**
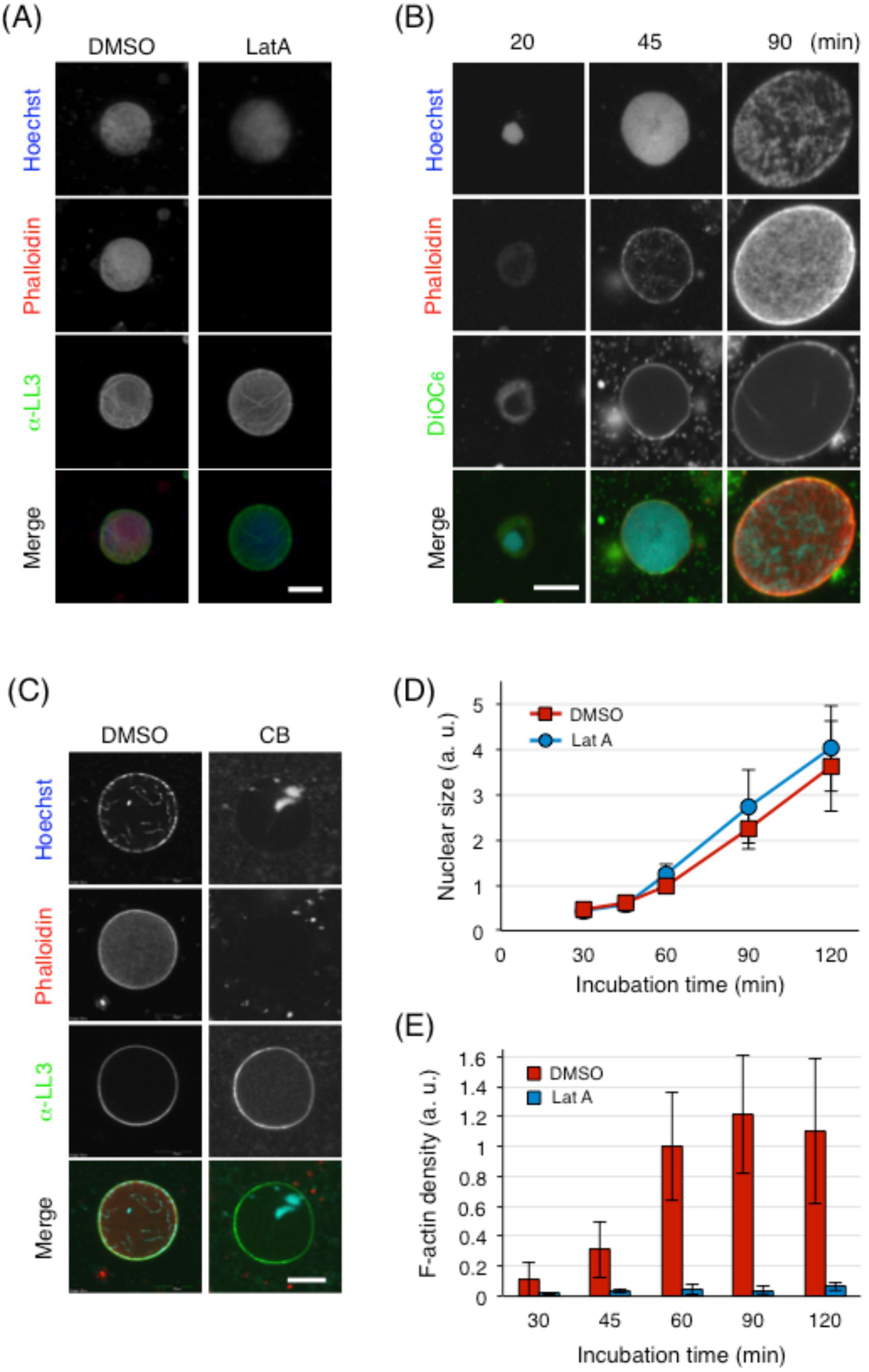
F-actin accumulates in nuclei formed in inhibitor-free egg extracts. (A) IFE(DMSO) and ICE (LatA: latrunculin A, 1.0 M) were supplemented with permeabilized sperm and were activated as described in Figure 2. Nuclei at 90 min after incubation were fixed; precipitated onto a cover slip; stained with Hoechst 33342, DY-590-phalloidin, and anti-lamin L3 (LL3); and observed by wide-field fluorescence microscopy. Scale bar, 20 m. (B) IFE was supplemented with permeabilized sperm and was activated as described above. Nuclei at the indicated time points after activation were fixed; stained with Hoechst 33342, DY-590-phalloidin, and DiOC_6_; and observed by confocal microscopy. Scale bar, 10 m. (C)Permeabilized sperm were added to IFE supplemented with 10 g/ml cytochalasin B (CB) or solvent (DMSO), followed by activation of IFE. Nuclei at 120 min after activation were fixed and stained with Hoechst 33342, anti-lamin L3 antibody, and DY-590-phalloidin for confocal microscopy. Scale bar, 20 m. (D) Change in the nuclear size as a function of incubation time. Nuclear size was obtained by measuring the nuclear area stained with anti-LL3 antibody on images taken by fluorescence microscopy (Figure S1). Red squares and blue circles represent nuclei formed in IFE (DMSO) and ICE (Lat A, 1.0 M), respectively. Means from two independent experiments are shown as relative values (DMSO, 60 min = 1). Error bars represent SD (n = 50). (E) Change in the nuclear F-actin density during the nuclear growth period. Nuclear F-actin intensity was obtained by measuring the nuclear DY-590-phalloidin signal on images taken by fluorescence microscopy (Figure S1). Nuclear F-actin density was calculated by dividing the nuclear phalloidin signal intensity by the nuclear area that had been stained with anti-LL3 antibody on fluorescence images with original threshold using NIS-element BR software (Nikon). Red and blue bars represent nuclei formed in IFE (DMSO) and ICE (Lat A, 1.0 M), respectively. Means from two independent experiments are shown as relative values (DMSO, 60 min = 1). Error bars represent SD (n = 50).

To investigate how F-actin accumulates in the nucleus, nuclei formed in IFE were fixed and fluorescently stained for confocal microscopy. In the course of nuclear assembly and growth, F-actin was first detectable and densely accumulated at the periphery of nuclei, including the nuclear lamina, and was also distributed uniformly throughout the nucleoplasm, occasionally forming a filamentous meshwork (Fig. 3B); F-actin that accumulates at the nuclear periphery and throughout the nucleoplasm are referred to as nuclear lamina F-actin and nucleoplasmic F-actin, respectively. This observation unambiguously demonstrates that F-actin accumulates in the nucleus, particularly at the nuclear lamina (Fig. 3C). Quantitative measurements of the phalloidin signal detected by wide-field microscopy (Figs 3A, S1 in Supporting Information) revealed that the density of nuclear F-actin increased along with nuclear growth, plateauing after 60 min while nuclei continued to grow (Fig. 3D, E). These results indicate that actin polymerization originates in the sub-nuclear membrane region, suggesting a distribution of actin nucleators in the inner nuclear membrane and/or the nuclear lamina.

### Nuclear F-actin protects chromatin from precocious aggregation

During the course of microscopic observation of nuclei formed in IFE (IFE-nuclei) and ICE(ICE-nuclei), we noticed that chromatin distributions in the nucleus were markedly different between the two. In ICE-nuclei, chromatin was initially distributed throughout the nucleus but formed aggregates, detaching from the nuclear envelope (NE) within 3 h of nuclear formation (Fig. 2B, C, CB). In contrast, chromatin remained distributed throughout the nuclei, slightly condensing as at prometaphase, following 3 h of incubation in IFE (Fig. 2B, C, DMSO). These observations suggest that nuclear F-actin may be involved in maintaining the binding of chromatin tothe NE, thereby preventing chromatin from condensing and/or aggregating precociously. To examine this possibility, we induced nuclear formation in ICE containing various concentrations of AI (CB or LatA) and observed nuclei by confocal microscopy after staining F-actin and DNA with phalloidin and Hoechst, respectively. When nuclei were formed in ICE containing very low concentrations of AI, F-actin accumulated in the nucleus as did in IFE-nuclei (Fig. 4A, 0.1 g/ml CB; Fig. 4B, 0.1 M LatA). In the nucleus, chromatin was mostly distributed in the sub-nuclear membrane region, delineating the nuclear contour, and was partially dispersed and somewhat condensed in the nucleoplasm (Fig. 4A, B), indicating chromatin binding to the NE. In contrast, chromatin formed aggregates, mostly detached from the NE, in nuclei formed in ICE containing high concentrations of AI that induced the disappearance of nuclear F-actin (Fig. 4A, 10 g/ml CB; Fig. 4B, 1.0 M LatA). In nuclei formed in ICE containing intermediate concentrations of AI, chromatin tended to form aggregates, and its distribution in the sub-nuclear membrane region was inconspicuous (Fig.4A, 1.0 g/ml CB; Fig. 4B, 0.3 M LatA); the thick actin bundles observed in these nuclei could be artifacts caused by fixation or incubation for long periods of time under confocal microscopy, as they were not discernible in observations of freshly fixed nuclei by wide-field fluorescence microscopy (Figs 1C and 2C, D, 3A). Thus, in nuclei that are assembled and grown in IFE, chromatin dissociates from the NE and forms aggregates, depending on the reduction in nuclearF-actin density.

**Figure 4.**
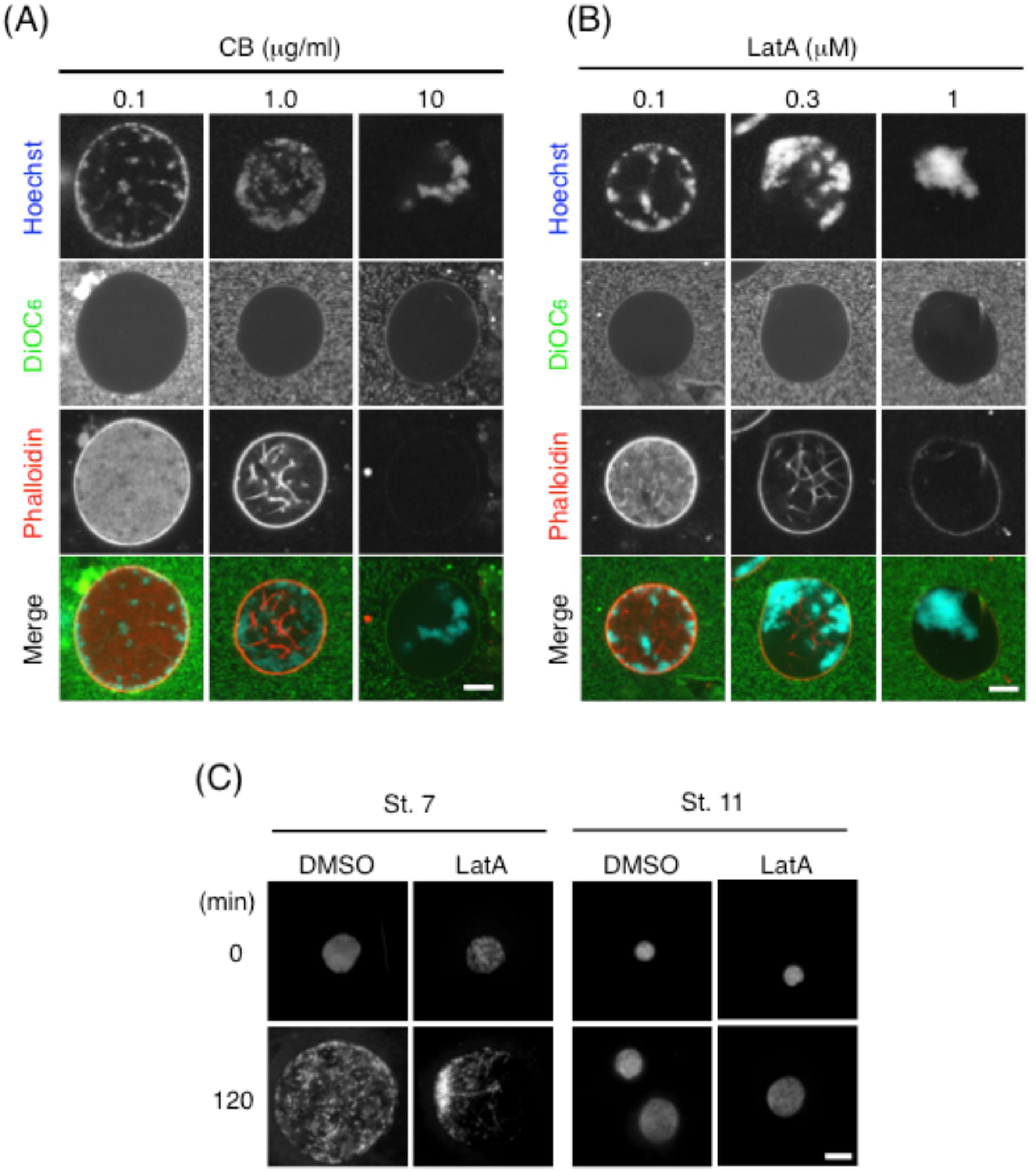
Nuclear F-actin maintains chromatin binding to the nuclear envelope in well-grownnuclei. (A and B) Permeabilized sperm were added to ICE supplemented with various concentrations of cytochalasin B (CB: 0.1, 1.0, or 10 g/ml) (A) or latrunculin A (LatA: 0.1, 0.3, or 1 M) (B) and incubated for 120 min. Nuclei were then fixed; stained with Hoechst 33342, DiOC_6_, and DY-590-phalloidin; and observed by confocal microscopy. Scale bars, 10 m. (C) Nuclei isolated from embryos at stage 7 (blastula) and stage 11 (gastrula) were added to IFE supplemented with latrunculin A (LatA: 1 M) or solvent (DMSO) and incubated for 120 min. Nuclei were then fixed, stained with Hoechst 33342, and observed by wide-field fluorescence microscopy. Scale bar, 10 m.

Although the above results clearly demonstrate that the accumulation of nuclear F-actin is required for the maintenance of NE–chromatin binding, it is possible that this requirement is a property of nuclei formed in egg extracts with sperm chromatin as the substrate and that this is not true for blastula nuclei. To examine this possibility, we isolated nuclei from blastulae at stage 7 and gastrulae at stage 11 and incubated them in IFE or ICE for comparison. When incubated in IFE, blastula nuclei grew to a much larger size (~50 m in diameter, Fig. 4C) after 2 h of incubation, like sperm chromatin nuclei (Fig. 2B). In these nuclei, the chromatin, which had slightly condensed, was distributed throughout the nucleus, maintaining its NE binding. By contrast, when incubated in ICE, blastula nuclei grew as they did in IFE, but, as expected, the chromatin aggregated, mostly detaching from the NE. In contrast, when the gastrula nuclei, which were originally smaller than the blastula nuclei, were examined, they did not grow as much as the blastula nuclei; interestingly, the distribution of chromatin in the nuclei remained unchanged after 2 h of incubation, regardless of whether they were incubated in IFE or ICE. These results strongly suggest that the necessity of nuclear F-actin for the maintenance of NE–chromatin binding is a character unique to the blastula nucleus. We therefore conclude that during the blastula stage, nuclear F-actin operates to maintain NE–chromatin binding and, presumably via this action, prevents chromatin from precociously aggregating in the nucleus.

### Nuclear F-actin stiffens the nuclear lamina

We anticipated that the observed increase in nuclear F-actin density in both the nuclear lamina and the nucleoplasm would stiffen the nucleus. We therefore examined the effect of nuclear F-actin accumulation on the structural stability of the nucleus. Egg extracts containing well-grown nuclei were diluted with the buffer used for preparing IFE, followed by high-speed centrifugation to precipitate the nuclei onto a coverslip. Precipitated nuclei were then fixed and fluorescently stained with an anti-lamin L3 (LL3) antibody and Hoechst for visualization of the nuclear lamina and chromatin, respectively (Fig. 5A). The resulting images showed that most IFE-nuclei maintained an intact nuclear lamina with chromatin distributed throughout the nucleus after centrifugation (Fig. 5A, B, DMSO), whereas more than half of ICE-nuclei contained a ruptured nuclear lamina with aggregated chromatin (Fig. 5A, B, LatA). It should be noted that numerous laminar fragments derived from broken nuclei were observed among ICE-nuclei, but these were rarely found among IFE-nuclei. Hence, in the case of ICE-nuclei, the proportion of broken nuclei was actually much higher than that presented in Fig. 5C. Thus, the structural stability of the nuclear lamina is increased by nuclear F-actin accumulation, likely because laminar F-actin directly increases stiffness by lining the nuclear lamina. In addition, nucleoplasmic F-actin should increase the viscosity of the nucleoplasm, rendering the nucleus more refractory to mechanical forces from the outside. It is also likely that nucleoplasmic F-actin indirectly supports the nuclear lamina through maintenance of NE–chromatin binding; because the nuclear lamina tends to rupture in chromatin-free regions (Fig. 5A, asterisks; Fig. 5B), the chromatin-bound nuclear lamina appears to be more stable than the unbound nuclear lamina.

**Figure 5.**
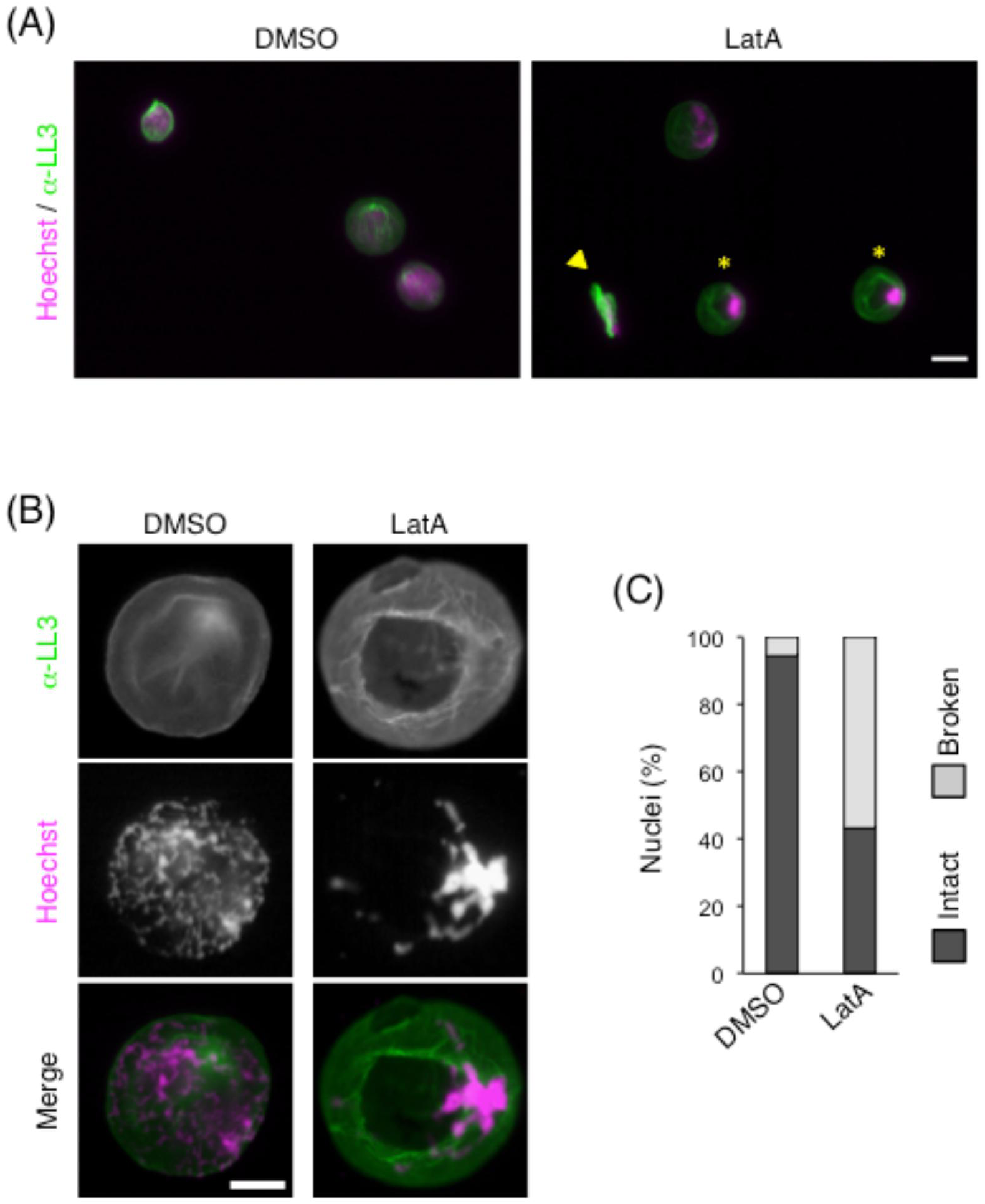
Nuclear F-actin stiffens the nuclear lamina in well-grown nuclei. (A) Permeabilized sperm were added to IFE supplemented with latrunculin A (LatA: 1.0 M) or solvent (DMSO) and incubated for 120 min. After extracts were diluted with extraction buffer, nuclei were precipitated onto a coverslip by centrifugation, followed by fixation and staining with anti-LL3 antibody and Hoechst 33342 and observation by wide-field fluorescence microscopy. Asterisks and arrowhead indicate nuclei with ruptured lamina and a laminar fragment, respectively. Scale bar, 20 m. (B) Representative images of each group are shown. Scale bar, 10 m. (C) Ratio of nuclei broken through centrifugal precipitation. Numbers of intact and broken nuclei in the experiment described in (A) were counted. Nuclei with intact and ruptured nuclear lamina (see DMSO and LatA in B) were regarded as intact and broken nuclei, respectively. Bar graph shows the percentage of intact and broken nuclei formed in IFE supplemented with DMSO (n = 195) or LatA (n = 914) after centrifugal precipitation.

### Nuclear F-actin facilitates chromosome alignment

In determining the physiological roles of nuclear F-actin, we also examined whether embryonic nuclear F-actin is involved in the regulation of chromosomal behavior during mitosis, arole thathas been suggested based on previous studies of nuclear F-actin in starfish oocytes(Lénárt *et al.* 2005; Mori **et al.** 2011). To investigate this, we induced nuclear formation in either interphase IFEor ICE and then transferred the nuclei to metaphase-arrested (M-) IFE to induce spindle assembly and chromosome alignment (Fig. 6A). When nuclei were incubated in M-IFE, they formed bipolar spindles on which condensed chromosomes were aligned (Fig. 6B), as has been reported for ICE (Sawin & Mitchison, 1991). Interestingly, mitotic spindles formed in IFE were often accompanied by F-actin masses (Fig. 6B, see also Fig. 1A, blastula), suggesting interactions between cytoplasmic F-actin and spindle microtubules. Measurements of the width and the length of the spindles indicated that there was no significant difference in spindle size between IFE-and ICE-nuclei (Fig. 6C, F/Fvs. -/F). Chromosome alignment, however, could be influenced by the presence or absence of nuclear F-actin in the earlier nuclei, namely whether the chromatin had precociously aggregated or not. When condensed chromosomes have been properly aligned on the metaphase plate of the spindle, the extent of chromosome scattering along the spindle axis is expected to be small. We therefore assessed the chromosome distribution on the spindles, measuring its length along the spindle axis and the width along the metaphase plate to calculate the ratio of the former to the latter; better chromosome alignment corresponds with a smaller ratio (Fig. 6D). Results indicated that chromosome alignment was significantly better in IFE-nuclei than in ICE-nuclei (Fig. 6D, F/F vs. -/F), suggesting that nuclear F-actin contributes to chromosome alignment in the subsequent M phase.

**Figure 6.**
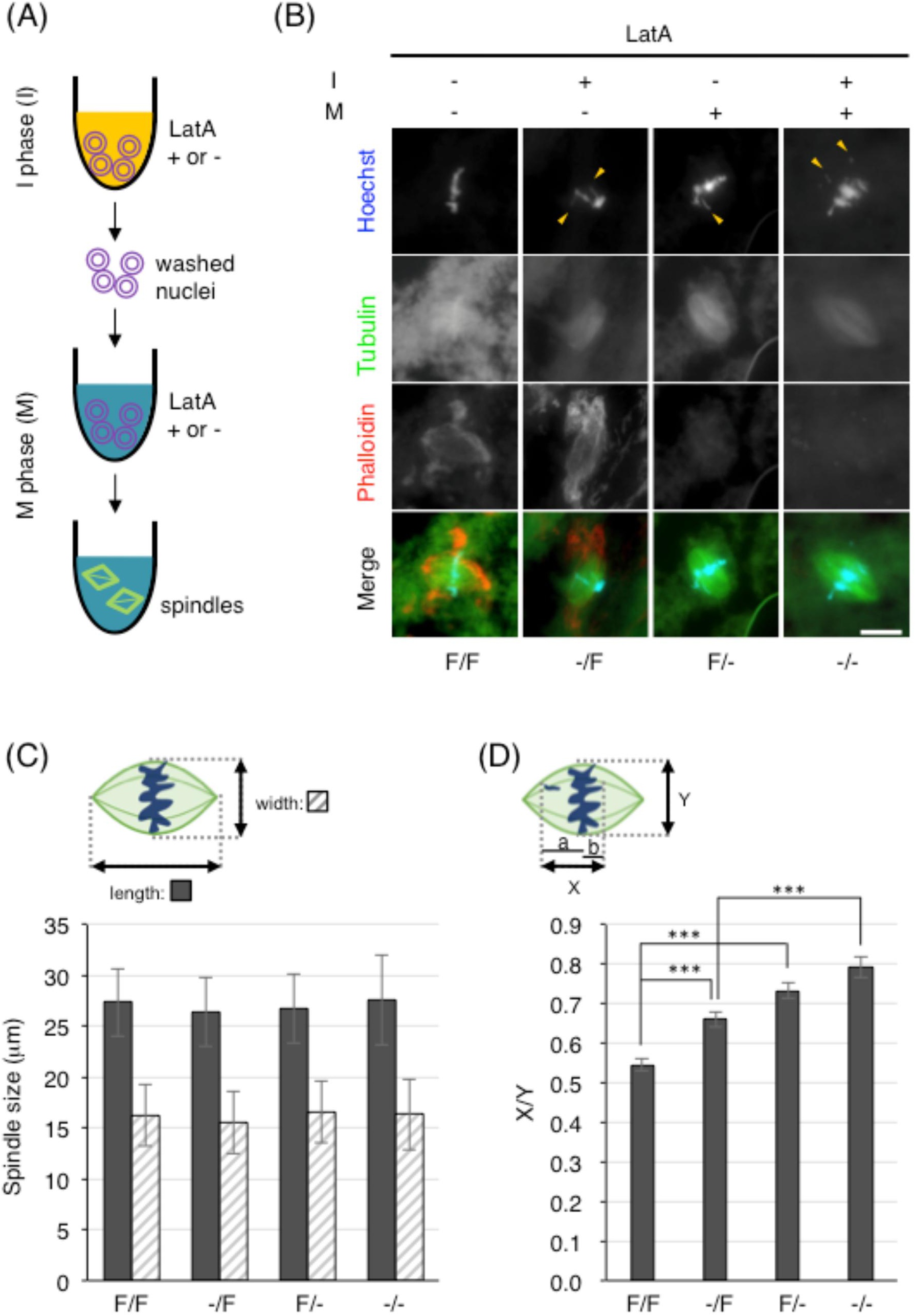
Nuclear F-actin facilitates chromosome alignment on the spindle. (A) Schematic of experiment to induce chromosome alignment on the spindle using egg extracts. Permeabilized sperm were incubated for 120 min in interphase IFE (I) supplemented with latrunculin A (LatA+) or DMSO (LatA -) to induce nuclear formation. After washing with extraction buffer, nuclei were incubated in M phase-arrested IFE (M) supplemented with latrunculin A (LatA +) or DMSO (LatA -) along with HiLyte Fluor 488-tubulin to induce spindle assembly. (B) Mitotic spindles assembled in M phase-arrested extracts were fixed and stained with Hoechst 33342 and DY-590-phalloidin and were observed by wide-field fluorescence microscopy. Presence (+) or absence of LatA (-) in interphase (I) and M phase-arrested (M) extracts is indicated at the top of the panel, and presence (F) or absence of F-actin (-) in egg extracts (I phase/M phase) is indicated at the bottom. Representative images of each experimental group are shown. Arrowheads indicate misaligned chromosomes. Scale bar, 20 m. (C) Spindle sizes. Lengths (black bars) and widths (hatched bars) of at least 150 spindles were measured for each of the four groups in (B). Bars and error bars represent means and SD, respectively. (D) Chromosome distribution area on spindles. The width (Y) and the length of chromosome distribution area (X = a + b) were measured on each of the spindles examined in (C). The extent of chromosome alignment is expressed as the ratio of X to Y (X/Y). Bars and error bars represent means and SE, respectively. **P < 0.001.

An association between F-actin masses and mitotic spindles may imply that F-actin is involved in the regulation of spindle formation and chromosome alignment. To examine this possibility, we compared spindles that were formed in M-IFE and M-ICE. The result clearly demonstrated that there was no difference in the sizes of spindles formed in M-IFE and M-ICE (Figure 6C, F/F vs. F/-, -/F vs. -/-). However, chromosome alignment was significantly better in M-IFE than in M-ICE, regardless of the presence or absence of nuclear F-actin in the preceding interphase (Fig. 6D, F/F vs. F/-, -/F vs. -/-). Taken together, these results suggest that in *Xenopus* blastomeres, F-actin contributes to chromosome alignment via spindle interactions during M phase and via nuclear accumulation during the preceding interphase, which prevents precocious chromatin aggregation.

## Discussion

In the present study, we demonstrated that F-actin accumulates in the nucleus during the blastula stage of *Xenopus* embryogenesis. Regarding nuclear F-actin in *Xenopus* embryos, it has been suggested that Exp6 expression during oocyte maturation should reduce the nuclear actin concentration in fertilized eggs, thereby making it unlikely for F-actin to accumulate in the nuclei of embryonic cells as it does in oocytes (Bohnsack *et al.* 2006). However, this study also demonstrated that expression of Exp6 increases gradually throughout the progression of embryogenesis, indicating that Exp6 expression in the blastula is relatively lower than that in the gastrula. Thus, during the blastula stage, a lower level of Exp6 allows for nuclear accumulation of actin monomers, as demonstrated by the use of cell-free extracts from *Xenopus* eggs (Krauss *et al.* 2003), at concentrations sufficient for actin polymerization. A subsequent increase in the Exp6 expression level should further decrease the nuclear actin concentration, to which the disappearance of nuclear F-actin at the gastrula stage may be at least partly attributable. In addition, during the cleavage stage, cell divisions with no growth occur at short intervals, bringing about a rapid, exponential increase in the cell surface area, which doubles every 90 minutes. Since F-actin accumulates at the sub-membranous region to organize the cell cortex, cleavage should result in a decrease in the cytoplasmic actin concentration and a corresponding decrease in the nuclear actin concentration. Thus, the nuclear actin concentration is thought to decrease gradually with the progression of embryogenesis. We have shown, however, that nuclear F-actin disappears abruptly at the gastrula stage in a nearly all-or-none manner. This suggests the involvement of a regulatory mechanism for nuclear F-actin, likely involving actin nucleators, independent of that responsible for the nuclear actin concentration.

We were surprised to find that nuclear F-actin is essential for the maintenance of NE–chromatin binding in the *Xenopus* blastula; when actin polymerization is inhibited with AI, chromatin detaches from the NE, forming aggregates in well-grown blastula nuclei. To our knowledge, this is the first evidence that nuclear F-actin is involved in the regulation of NE– chromatin binding. In terms of the mechanism by which F-actin tethers chromatin to the NE, it is unlikely that F-actin interacts directly with chromatin, as no physical interactions between F-actin and chromatin were observed through confocal imaging. Rather, we emphasize that nuclear F-actin first becomes detectable and densely accumulates at the nuclear lamina. This observation implies that within the nucleus, actin nucleators are distributed in the nuclear lamina region similar to the way in which they are localized to the cell cortex. Whatever the mechanism, F-actin densely accumulates at the nuclear lamina to form a layer of proteinaceous matrices in the sub-inner nuclear membrane region, as F-actin does in the cell cortex. The chromatin, which is associated with the NE during nuclear assembly and growth, should be embedded, at least in part, in these actin matrices. We speculate that nuclear F-actin thus tethers chromatin to the NE. According to this hypothesis, the machinery that mediates NE–chromatin binding upon nuclear assembly and during the early stages of nuclear growth should be unable to protect the chromatin from dissociating from the NE during the rapid expansion of the NE that occurs in the late stages of nuclear growth. In this regard, we note that the inner nuclear membrane protein (INMP) composition in *Xenopus* eggs and blastomeres is unique and different from that of differentiated cells, including gastrula cells (Lang *et al.* 1999; Gareiß *et al.* 2005). New evidence suggests that in nuclei formed in egg extracts, the chromatin tends to detach from the NE owing to the uniqueness of the INMP composition (manuscript in preparation). Differences in the INMP and lamin compositions of blastula and gastrula nuclei(Benavente *et al.* 1985; Stick & Hausen, 1985) explain their distinct changes in egg extracts. The unique INMP composition in *Xenopus* eggs and blastomeres may also be related to the blastula-specific nuclear F-actin accumulation.

In this study, we provided evidence that nuclear F-actin increases the structural stability of the nucleus and facilitates chromosome alignment on the spindle in the following M phase. During the cleavage period, nuclear formation and breakdown alternates at short intervals, and mitoses are undertaken in the cytoplasms of cleaving blastomeres, which should be moving dynamically owing to cytokinesis of the large cells. Under these circumstances, both contributions of nuclear F-actin are likely to be beneficial in maintaining chromosomal integrity. Notably, similar contributions of nuclear F-actin have been demonstrated in the GV of *Xenopus* (Bohnsack *et al.* 2006) and in starfish oocytes (Lénárt *et al.* 2005).

In *Xenopus* embryogenesis, zygotic transition first becomes detectable in the mid-blastula at stage 8 concurrent with the mid-blastula transition (MBT), and thereafter, transcription levels gradually increase to a plateau in the gastrula at around stage 11 (Newport & Kirschner, 1982). Thus, in *Xenopus* embryos, zygotic transcription is fully activated during the transition period from the late blastula (stage 9) to the early gastrula (stage 10). It should be underscored that transcriptional activation occurs concomitantly with the disappearance of nuclear F-actin. Correspondingly, it is intriguing that a recent study has shown that mammalian somatic nuclei can be transcriptionally reprogrammed through transplantation of *Xenopus* oocytes into the GV and that nuclear actin polymerization is required for the induction of the reprogramming (Miyamoto *et al.* 2011). Taken together, these findings imply that blastula-specific nuclear F-actin accumulation may be related to the induction and/or maintenance of the initialized, undifferentiated state of the nucleus in *Xenopus* eggs and blastomeres.

In summary, we demonstrated that in the *Xenopus* blastula, F-actin accumulates in the nucleus and maintains NE–chromatin binding. Although the mechanism of nuclear F-actin accumulation remains to be elucidated, it is clear that nuclear F-actin increases the structural stability of the nucleus and facilitates chromosome alignment on the spindle, thereby contributing to the maintenance of chromosomal integrity in rapidly dividing, large *Xenopus* blastomeres. Since the nucleoskeletal function of F-actin in early *Xenopus* embryos is likely related to the cellular properties of blastomeres, this may be common among animal blastomeres that undergo cleavage divisions at short intervals. Considering that evidence for the involvement of nuclear F-actin in chromatin regulation has been accumulating recently, the regulatory mechanism of nuclear F-actin accumulation and its relevance to transcriptional control in *Xenopus* early embryos deserve future study.

## Experimental procedures

### *Xenopus* embryos, microinjections, and nuclear preparation

*Xenopus* embryos were obtained by artificial insemination, de-jellied in 1.25% thioglycolic acid (pH 8.2), and allowed to develop in 10% MMR (100 mM NaCl_2_, 2 mM KCl, 1 mM MgCl_2_, 2 mM CaCl_2_, 0.1 mM EDTA, 5 mM HEPES-NaOH, pH 7.8). Some de-jellied embryos were microinjected with recombinant proteins and mRNAs during the one-cell stage and incubated in 10% MMR containing 5% Ficoll. Prior to nuclear preparation, embryos were treated with cycloheximide (50 g/ml) for 30 min to increase interphase cells. Embryos were staged according to Nieuwkoop and Faber (1967).

For nuclear preparation, 15 de-jellied embryos were washed in ice-cold sucrose extraction buffer (SEB: 250 mM sucrose, 100 mM KCl, 5 mM MgCl_2_, 20 mM HEPES-KOH, pH 7.4), suspended in 60 l SEB, and broken by passing through a wide-orifice tip (QSP, 118-N-Q) several times. Embryos were further broken by passing through a wide-orifice tip that had made contact with the inside wall of the tube bottom. For efficient nuclear preparation, early, mid, and late blastulae and gastrulae were passed through the tip 1–3, 5–10, and 15–20 times, respectively. Suspensions of broken embryos were immediately mixed with an equal volume of 4% paraformaldehyde (PA)-PBS and incubated at room temperature for 5 min.

### Nuclear formation in egg extracts

Inhibitor-free extracts (IFE) were prepared from unfertilized *X. laevis* eggs as previously described (Yamamoto *et al.* 2005) except that the EGTA-extraction buffer (EEB: 100 mM KCl, 5 mM MgCl_2_, 0.1 mM CaCl_2_, 5 mM EGTA, 20 mM HEPES-KOH, pH 7.4) was not supplemented with cytochalasin B. Briefly, unfertilized *Xenopus* eggs were de-jellied with 2.5% thioglycolic acid-NaOH (pH 8.2), washed five times with EEB, and packed into a plastic tube. After removal of excess buffer, eggs were centrifuged at 15,000 *g* for 10 min at 4°C. The cytoplasmic fraction between the lipid cap and precipitated yolk was removed and centrifuged again at 15,000 *g* for 15 min at 4°C. After the second centrifugation, the cytoplasmic fraction between the lipid and membranous precipitates was removed (Figure S2) and homogenized by pipetting before use. Thus-prepared egg extracts were arrested at metaphase (M-phase extract). To induce nuclear formation, M-phase extracts were supplemented with membrane-permeabilized sperm nuclei (750 nuclei/l; Ohsumi *et al.* 2006) and cycloheximide (100 g/ml) and were then activated by adding 0.4 mM CaCl_2_ to induce cell-cycle transition to S phase (Murray, 1991).

### Fluorescence microscopic observations of nuclei

For wide-field fluorescence microscopic observation, 2 l of PA-fixed embryo suspensions were mixed with 1.5 l of a nuclear staining solution (NSS: 5 g/ml Hoechst 33342, 20 g/ml DiOC_6_, 30% glycerol, and 10% formalin in EB: 100 mM KCl, 5 mM MgCl_2_, 20 mM HEPES-KOH, pH 7.4; Ohsumi *et al.* 2006) and 0.5 l of a DY-590-phalloidin solution (200 units/ml). For observation of nuclei formed in egg extracts (extract nuclei), 2 l egg extracts containing nuclei were mixed with an equal volume of NSS. Nuclei were visualized with a Nikon Eclipse 80i fluorescent microscope, and images were acquired with a digital CCD camera (Nikon DS-2MBWc).

For confocal microscopic observation of extract nuclei, egg extracts containing nuclei were diluted 1:10 with 4% PA-PBS and incubated at 22°C for 5 min and then at 4°C overnight. Fixed extracts were supplemented with DY-590-phalloidin (0.4 units/ml), Hoechst 33342 (2 g/ml), and DiOC_6_ (10 g/ml); incubated for 10 min; and layered over a step gradient cushion (250 l of 30% sucrose-EB on 50 l of 2 M sucrose-EB). This was followed by centrifugation for 10 min at 22°C at centrifugal forces ranging from 800 to 2,000 *g* depending on the extent of nuclear growth. After centrifugation, a 20-l fraction on the 2 M sucrose cushion was transferred to a single-well glass-base dish filled with mineral oil. Confocal images were acquired with an Olympus FV10i-DOC.

In some experiments, egg extracts were supplemented with His-EGFP-UtrCH (50 g/ml) before sperm addition, and nuclei were observed immediately after the addition of the staining solution by wide-field fluorescence microscopy. For confocal microscopy of unfixed nuclei that accumulated EGFP-UtrCH protein, egg extracts were diluted 1:10 with EB and processed as described above for fixed nuclei except that nuclei were precipitated by centrifugation at 1,000 *g* for 10 min at 4°C.

### Immunofluorescent staining of extract nuclei

For wide-field fluorescence microscopic observation, immunofluorescent staining of extract nuclei was performed according to the method described by Iwabuchi *et al.* (2002) with modifications. Briefly, 10–20 l of egg extracts containing nuclei were mixed with 1 ml of 4% PA-PBS and incubated for 2 h at room temperature. Fixed extracts were layered over a 1-ml cushion (30% sucrose in EB), and nuclei were precipitated onto a polylysine-coated coverslip through the sucrose cushion by centrifugation at 2,000 *g* for 5 min at 22°C. After treatment with 0.1% Triton X-100 in PBS for 1 min at room temperature, coverslips were washed three times in PBS and blocked in 2% BSA-PBS solution. Coverslips were then incubated with anti-*Xenopus* lamin L3 (LL3) antibody (1:500 dilution; Hasebe *et al.* 2011), followed by incubation with Alexa 488-conjugated secondary antibody (1:1000 dilution; Invitrogen) and DY-590-phalloidin (2 units/ml, Dyomics GmbH). Coverslips were washed in PBS, treated with Hoechst 33342 (2 g/ml) and mounted on glass slides with a mounting reagent (SlowFade Gold, Molecular Probes). In some experiments, extract nuclei were precipitated for immunostaining onto a coverslip prior to fixation, as follows. Egg extracts (20 l) containing nuclei were diluted with 1 ml SEB and layered over a 1-ml cushion of 30% sucrose-EB. Nuclei were precipitated onto a polylysine-coated coverslip through the sucrose cushion by centrifugation at 2,430 *g* for min. Coverslips were then treated with 4% PA-PBS for 5 min at 22°C and processed for immunofluorescence staining with anti-LL3 antibody and counterstaining with Hoechst 33342.

For confocal microscopic observation, PA-fixed extracts were supplemented with 025% Triton X-100, incubated at 22°C for 5 min, and layered over the step gradient cushion (250 l of 30% sucrose-EB on 50 l of 2 M sucrose-EB). After centrifugation (8,000 *g*, 5 min, 22°C), a 200-l fraction from the tube bottom, which included the 2 M-sucrose step and a part of the 30% sucrose step containing nuclei, was removed and diluted with 1.0 ml of 2% BSA-EB, followed by incubation for 30 min at 22°C with gentle agitation. For the primary antibody reaction, anti-LL3 antibody was added to the nuclear suspension and incubated for 1 h at 22°C with gentle agitation. The nuclear suspension was layered over a 50 l cushion of 2 M sucrose-EB and centrifuged at 5,000 *g* for 10 min at 22°C. A 200-l fraction from the tube bottom was removed, diluted as described above, and then incubated with Alexa 488-conjugated secondary antibody (1:1000 dilution), DY-590-phalloidin (0.4 units/ml), and Hoechst 33342 (2 g/ml) for 1 h at 22°C with gentle agitation. The nuclear suspension was layered over a 50-l cushion of 2 M sucrose-EB, and, after centrifugation (5,000 *g*, 10 min, 22°C), a 20-l fraction from the 2 M-sucrose cushion was transferred to a single-well glass-base dish for confocal microscopy as described above.

### Nuclear F-actin density

To quantitate nuclear F-actin density, PA-fixed embryo suspensions were stained with NSS and DY-590-phalloidin as described above. For extract nuclei, egg extracts containing nuclei were diluted 1:9 with 4% PA-PBS for 2 h, followed by precipitation onto coverslips, immunofluorescent staining with anti-LL3 antibody, and staining with DY-590-phalloidin and Hoechst 33342 as described above. The intensity of nuclear phalloidin signal and the nuclear area stained with DiOC_6_ (embryo nuclei) and anti-LL3 (extract nuclei) were quantified based on images obtained by wide-field fluorescence microscopy as described above.

### Mitotic spindle assembly

Egg extracts (25 l) containing nuclei at 120 min after activation were diluted to 1 ml with SEB, layered over a 400 l-cushion (30% sucrose-EB), and centrifuged at 2,000 *g* for 3 min at 4°C. After washing twice with 1 ml SEB, precipitated nuclei were incubated in M-phase extracts supplemented with HiLyte Fluor 488-labeled tubulin (cytoskeleton, 0.08 g/l) for 90 min at 22°C. For both interphase and M-phase extracts, those supplemented with latrunculin A (1 µM) or solvent (DMSO) were used. Extracts were mounted on glass slides with an equal volume of mounting reagent containing 10% formalin, Hoechst 33342 (5 g/ml), and DY-590-phalloidin (2 units/ml) and observed by wide-field fluorescent microscopy.

### Recombinant proteins

The EGFP gene was cloned into pET30b to obtain the pET30b-EGFP plasmid. A cDNA fragment encoding the F-actin binding domain (1–261, calponin homology domain) of human utrophin (UtrCH) was cloned into the pET30b-EGFP plasmid. His_6_-EGFP-UtrCH was expressed in *E. coli* strain BL21 (DE3). Cells were grown in 2× YTG medium supplemented with kanamycin (50 g/ml) at 37°C overnight. Cultures were diluted ten-fold into 2× YT and shaken at 18°C to an OD_600_ of 0.6. After addition of 0.2 mM IPTG, cells were shaken for 20 h at 18°C to express the recombinant protein and then kept on ice for 90 min. Harvested cells were lysed by sonication, and His_6_-EGFP-UtrCH was purified from the soluble fraction using His·bind resin (Novagen). After elution, His_6_-EGFP-UtrCH was dialyzed against EB, concentrated to 5 mg/ml, and stored at −80°C.

### mRNA synthesis

EGFP-UtrCH261, Lifeact-2EGFP, and XH2B-mTagBFP mRNA were synthesized *in vitro* using the mMESSAGE mMACHINE T3 kit (Ambion). mRNAs were dissolved in water and used for microinjection experiments.

### Plasmids

A cDNA fragment encoding the calponin homology domain (1–261) of human utrophin (UtrCH, NM_007124; Burkel *et al.* 2007) was cloned from RPE1 cells by RT-PCR. Two oligonucleotides were used as 5′ (5′-CAGTACTATGGCCAAGTATGGAGAACATGAAGCC-3′) and 3′ (5′- GCTCGAGGTCTATGGTGACTTGCTGAGGTAGCAC -3′) primers containing synthetic *Sca* I and *Xho* I restriction sites (underlined), respectively. PCR products were cloned into the pGEM-T Easy plasmid (Promega). A *Sca* I-*Xho* I fragment of the UtrCH gene was fused to the carboxy-terminus of the EGFP gene in the pET30b vector (Novagen) to generate pET30b-EGFP-UtrCH with His_6_ tags at the amino-and carboxy-termini of the fusion protein.

For *in vitro* mRNA synthesis, cDNAs were cloned into plasmids derived from pBS-RNT3 (Sawada *et al.*, 2005), which carries the UTRs of *X. laevis* globin mRNA (Lemaire *et al.* 1995). For construction of a plasmid to synthesize EGFP-UtrCH mRNA, the *UtrCH* gene was amplified by PCR using 5′ (5′- CGGATCCATGGCCAAGTATGGAGAACATGAAGCC-3′) and 3′ (5′- CTCGAGTTAGTCTATGGTGACTTGCTGAGGTAGC-3′) primers containing synthetic *Bam* HI site and *Xho* I site with a stop codon (underlined), respectively, and cloned into the pGEM-T Easy plasmid (Promega). A *Bam* HI-*Not* I fragment of the *UtrCH* gene was ligated to the pEGFP-N1 plasmid to obtain pEGFP-N1-UtrCH. For construction of a plasmid to synthesize Lifeact-2EGFP mRNA, synthetic oligonucleotides for sense (5′-ATCTATGGGAGTGGCTGATCTGATTAAGAAGTTTGAATCTATTTCTAAGGAAG AAGGAGGATCTGGAT-3′) and antisense (5′-CATGATCCAGATCCTCCTTCTTCCTTAGAAATAGATTCAAACTTCTTAATCAGATCAGCCACTCCCATA-3′) strands containing the Lifeact sequence (Riedl *et al.* 2008) were annealed and then ligated into the pEGFP-C4 plasmid digested with *Bgl* II and *Nco* I to generate pLifeact-EGFP. The *EGFP* gene was further ligated to the pLifeact-EGFP plasmid to generate pLifeact-2EGFP. A cDNA encoding histone H2B (NM_001093284) was isolated from *X. laevis* eggs by RT-PCR using 5′ (5′-GAGATCTATGCCTGAGCCCGCCAAATCCGCTCC-3′) and 3′ (5′- GCCATGGCTTGGCGCTGGTGTACTTGGTGACGG-3′) primers containing synthetic *Bgl* II and *Nco* I sites (underlined), respectively, and cloned into the pGEM-T Easy plasmid. The gene for *mTagBFP* (Subach *et al.* 2008) was synthesized with a flexible linker sequence (encoding 15 amino acids of the repeat Gly-Gly-Ser) fused to the amino-terminus. A *Nco* I-*Not* I fragment of (GGS)_5_-mTagBFP was fused to the carboxy-terminus of the *H2B* gene in the pBS-RNT3 plasmid to generate pBFP-C4-H2B.

## Acknowledgements

We thank M. Harata for helpful advice with confocal microscopy, R. Uehara for RPE1 cell cDNAs, the Nagoya University Center for Gene Research for the use of a confocal microscope, and K. Miyamoto and T. Nakayama for critical reading of the manuscript. This work was supported by JSPS KAKENHI Grant Numbers, JP26650005 (M. I.) and JP26650058 (K. O.).

## Author contributions

M. I. and K. O. conceived the project. K. O. and N. U. devised inhibitor-free egg extracts. H. O., N. S., and M. I. performed most of the experiments and analyzed the data. M. I. and K. O. designed the experiments and wrote the paper.

## Supporting information

**Figure S1.**
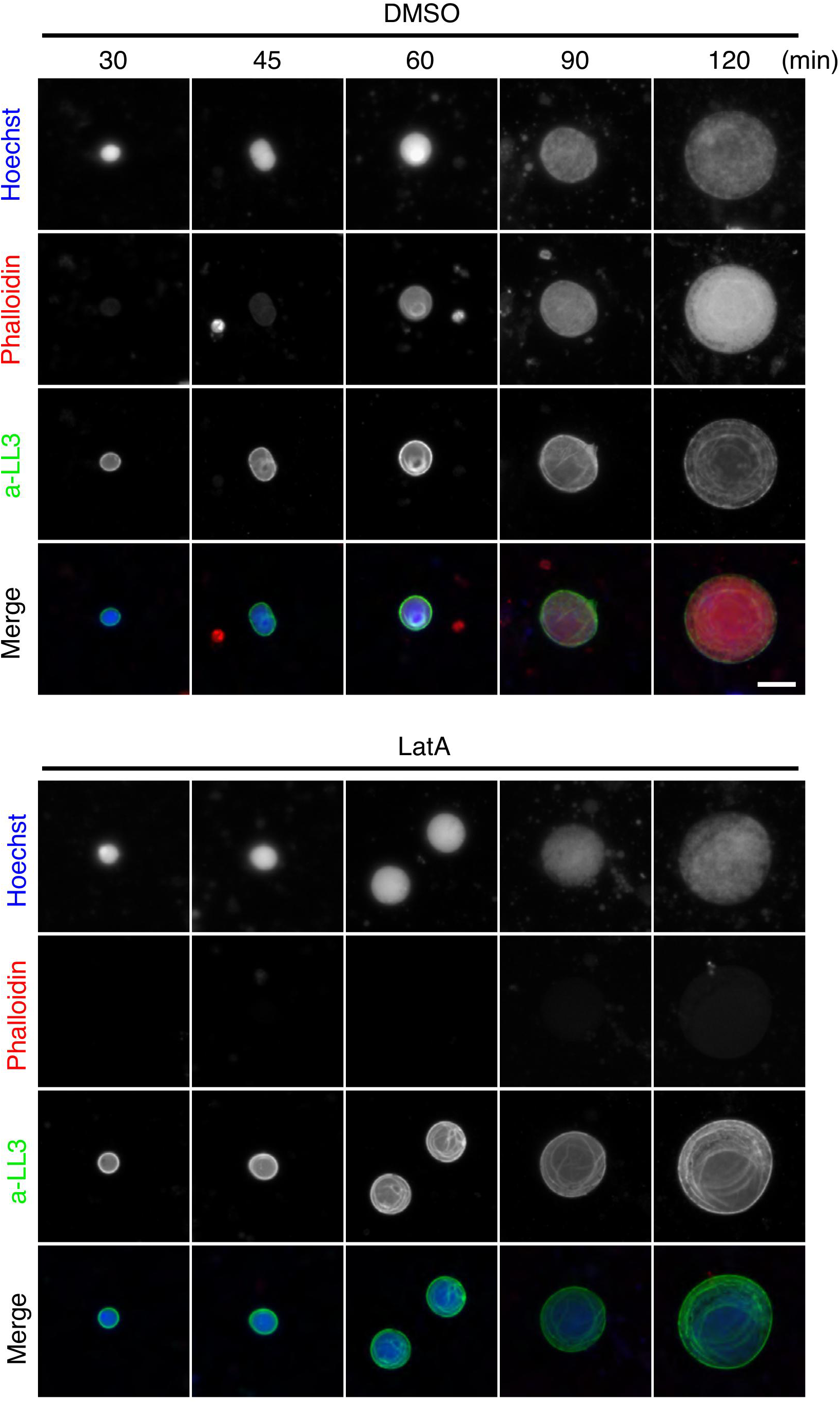
F-actin accumulates in nuclei formed in inhibitor-free egg extracts (related to Figure 2). Permeabilized sperm were added to IFE supplemented with 1.0 μM latrunculin A (LatA) or solvent (DMSO), followed by activation of IFE. Nuclei at the indicated time points after activation were fixed, precipitated onto a coverslip by centrifugation, and stained with Hoechst 33342, anti-lamin L3 antibody, and DY-590-phalloidin for wide-field fluorescence microscopy. Scale bar, 20 μm.

